# The toxic effects of lead nitrate on human keratinocytes (HaCaT)

**DOI:** 10.1101/2024.11.20.624611

**Authors:** Adnan Mohamed, Pemma Mahathi, Niveditha Nair, Herman DSouza

## Abstract

This study investigated the cytotoxicity, genotoxicity, oxidative stress, and mitochondrial toxicity in human keratinocytes (HaCaT) caused by lead nitrate. An increase in lead nitrate concentration (50-800 µM) showed a decrease in cell viability and the IC_50_ concentration was found to be 385.7 µM. Lead nitrate at 250 µM was also found to decrease the clonogenic potential, produce an increased number of binucleated cells with a micronucleus, increase the olive tail movement, reduce the thiol protein content, increase the intracellular and mitochondrial ROS levels, increase the mitochondrial mass and mitochondrial membrane potential. Which showed that Lead nitrate does indeed show cytotoxic, genotoxic effects as well as increase oxidative stress and alter mitochondrial functions.

**HIGHLIGHTS:**

- Lead nitrate causes cytotoxicity in keratinocytes with an IC_50_ conc. of 385.7 µM.
- Exposure to lead nitrate showed a significant increase in DNA damage.
- Lead nitrate increased the mitochondrial membrane potential and mass.

## INTRODUCTION

Lead, a metal of significant anthropogenic relevance and notable toxicity, has been recognized since ancient times. Historically considered by alchemists as one of the earliest known metals, lead is derived from the Latin term ‘plumbum,’ meaning ‘liquid silver.’ This bluish-grey element, with an atomic number of 82, has become one of the most pervasive metals utilized by humanity. It is detectable across all biological organisms and in diverse environmental conditions (Flora et al., 2012). Lead is extensively distributed worldwide, owing to its notable properties such as malleability, ductility, ease of welding, and resistance to corrosion. These characteristics have led to its widespread application across various industrial sectors, including plastics, ceramics, paints, and automobiles (Cotter-Howells and Thornton, 1991). Exposure to lead occurs through various routes, including ingestion, inhalation, and dermal contact. In the 1980s it was estimated that daily lead intake ranges from 150-300 µg via oral ingestion and 10-20 µg via the respiratory tract. This exposure significantly impacts basal physiological functions, with children being particularly vulnerable compared to adults (Murray, 2004). Recent studies show that children in the United States are exposed to 1.4–1.8 μg/day of lead just through their dietary intake, so there has been a notable reduction in the intake of this harmful metal.

Although lead toxicity affects almost all the organs, its impact on the skin remains largely understudied. Skin, often overlooked in the context of heavy metal toxicity, exhibits a range of adverse effects upon lead exposure, including compromised hydration, reduced elasticity, hyperpigmentation. The appearance of a “lead hue” on the skin, as well as different dermatological anomalies. Symptoms of skin damage is a simple technique to detect chronic lead toxicity in asymptomatic patients (Rerknimitr et al., 2019; Talaie et al., 2018).

The main mechanisms of lead toxicity involve the direct generation of ROS through fenton reactions and redox cycling or indirect oxidative stress caused by lipid peroxidation (Sharma et al., 2015). Lead possesses an affinity for sulfhydryl (-SH) groups, thereby interfering with the functioning of various enzymes containing these groups such as enzymes involved in antioxidant response and heme synthesis (Lakka et al., 2023). Lead is also known to act as calcium analogue which causes it to disrupt ion channels and vitamin D synthesis (Wani et al., 2015).

While acute exposure to high levels of lead can result in lead poisoning, characterized by extreme abdominal pain, vomiting, peripheral neuropathies, and encephalopathy, such high levels of exposure are now rare. This rarity is due to the extensive risk assessment and phase-out measures implemented by major countries, such as the U.S., since the 1970s. These measures have significantly reduced dependency on lead. Nevertheless, major sources of lead exposure persist, including house paints, corroded pipelines, automobile fuel, and gasoline. Occupational exposure remains a significant concern for workers in mining, smelting, and recycling plants, where lead levels are exceptionally high (Cookman et al., 1987).

Chronic lead exposure, which most people encounter over long periods, is responsible for numerous health deficits later in life. These include neurodegenerative disorders, cognitive and behavioural deficits, miscarriages in pregnant women, gestational hypertension, learning impairments in children, reduced IQ, and long-term dysfunction of several physiological systems such as neurological, reproductive, hematopoietic, and cardiovascular systems (Mason et al., 2014).

The estimation of blood lead levels can be conducted using K-X-Ray Fluorescence (Somerville et al., 1985), Atomic Absorption Spectroscopy (AAS) (Amiri and Amini, 2012), or Inductively Coupled Plasma Mass Spectrometry (ICP-MS), depending on the available resources (Canfield et al., 2003). Various sample types, including whole blood, plasma, serum, nails, or hair, can be used to estimate lead burden in the body. Between 95-99% of ingested lead is absorbed by erythrocytes and subsequently distributed into soft tissues and bones. This distribution is facilitated by lead’s ability to mimic calcium, allowing its transfer from blood to bone during calcium resorption processes. The residual half-life of lead in blood is approximately 30 days, after which it is excreted in urine. However, lead can remain stored in bone for up to 20 years (Charkiewicz & Backstrand, 2020).

## MATERIALS AND METHODS

### 1. Chemicals/reagents

Lead nitrate (Pb(NO_3_)_2_), Acridine orange (AO), Rhodamine123, 5,5′-dithiobis(2-nitrobenzoic)acid (DTNB), Low melting Agarose were purchased from Sigma Chemical Co.(St. Louis, MO, USA), Dulbecco’s Modified Eagle Medium (DMEM), phosphate buffered saline (PBS), Fetal Bovine Serum (FBS), Dimethyl sulfoxide (DMSO), crystal violet, cytochalasin-B, Colchicine, Trypsin, Ethidium bromide, Agarose, Ethylene diamine tetra acetic acid (EDTA), Trypan blue, Giemsa, 3-(4,5-dimethylthiazol-2-yl)-2,5-diphenyltetrazolium bromide (MTT), Hanks Balanced Salt Solution (HBSS), Potassium Chloride (KCl) were purchased from HiMedia Laboratories Pvt.Ltd., Nashik, India, 2′,7′-dichlorofluorescindiacetate (DCFDA), MitoSOX Red Dye, nonyl acridine orange (NAO) were purchased from ThermoFisher Scientific, USA.

### 2. Cell culture conditions

This work used HaCaT, a human keratinocyte cell line that had been spontaneously immortalized. T-25 flasks were used to grow them with DMEM supplemented with 10% FBS at 37°C in an incubator (NuAire, Plymouth, MN, USA) in an environment with humidified air with 5% CO2.

### 3. Preparation of Lead nitrate

10 mM solution of Pb(NO_3_)_2_ was made with milliQ water and was further diluted with DMEM as necessary for the study.

### 4. Analysis of cell viability by MTT assay

Cell viability after exposure to different concentrations of Pb(NO_3_)_2_ was ascertained using the MTT assay (Mosmann, 1983). HaCaT cells which were exponentially growing was trypsinised and 5000 cells were plated in each well in a 96 well and treated with Pb(NO_3_)_2_ conc. taken from 50 µM -800 µM. Then the treatment along with the media was removed and 100µl of 5µM MTT in HBSS was added, incubated for 4 hrs, then the MTT solution was removed and 100µl DMSO was added, and the absorbance was read at 570 nm using a plate reader (Infinite M200; Tecan, Austria).

### 5. Clonogenic assay

An appropriate number of cells was seeded onto petri dishes and were treated with 250 µM Pb(NO_3_)_2_ for 72 hrs. After the treatment the media is discarded and fresh DMEM was added, and the cultures are allowed to grow undisturbed for 10 days for colony formation in an incubator at 5% CO2 and 37°C (Puck & Marcus, 1955). The surviving fraction was then calculated by staining the dish with crystal violet and counting the colonies (Satish Rao et al., 2009).

### 6. Micronucleus assay

The genotoxicity induced by Pb(NO_3_)_2_ was assessed by the micronucleus assay.3×10^5^ cells were seeded into 6cm^2^ petri dishes and were treated with 250 µM Pb(NO_3_)_2_ for 72 hrs, followed by addition of 5 µM of Cytochalasin B for 24 hrs with fresh media. The cells are then trypsinised and collected in tubes, centrifuged at 1000 RPM for 10 min, the supernatant removed and washed with PBS, the tube is then vortexed followed by centrifugation, the supernatant was removed and then 5:1 Methanol and Acetic acid fixative was added while gently mixing the cells and stored at 4°C for 4 hrs. The cells were gently dropped onto a chilled slide washed with the fixative, and then stained with Acridine Orange and dipped in PBS, and viewed under a fluorescent microscope (Olympus CKX53, Tokyo, Japan). 10^3^ cells with well-preserved cytoplasm and a binucleated nucleus (BN) were scored from each of the cultures and the frequency of micronucleated binucleate cells (MNBNC) was analyzed (Fenech et al., 2003).

### 7. Comet assay

Comet assay or Alkaline single cell gel electrophoresis was conducted as mentioned elsewhere (Singh *et al*., 1988) and the DNA damage was assessed according to the change in olive tail moment (OTM). Cells were mixed with low melting point agarose and spread on a slide precoated with agarose, then it was submerged into a lysis buffer overnight to break down cell membrane and release DNA. The slides were then placed in an alkaline solution to unwind DNA which is followed by electrophoresis (16 V, 19 min) which creates the comet shape, the slide is then neutralized and stained with EtBr for visualization.

### 8. Evaluation of intracellular reactive oxygen species (ROS)

Treated cells were incubated with 5uM DCF-DA for 30 mins. Then the cells were washed, harvested, pelleted, and resuspended in PBS and was used to perform flow cytometry Cyflow® Space Partec, IL, USA) (excitation-488 nm; emission-525 nm) (Bai & Cederbaum, 2003).

### 9. Evaluation of intracellular protein thiol content

Intracellular protein thiol content was evaluated by 5,5′-dithiobis(2-nitrobenzoic) acid (DTNB). The treated cells were separated using trypsinization, lysed with cold trichloroacetic acid (6.5%), and pelleted. Then the pellet was combined with a reaction buffer with 100 *µ*M DTNB, 0.5 M Tris, 0.5% SDS, 0.5mM EDTA present. 2-nitro-5-thiobenzoate (TNB) is produced when DTNB and thiol groups react, and absorbance was measured at 412 nm (Rathi et al., 2017).

### 10. Evaluation of mitochondrial membrane potential (ΔΨm)

Treated cells were incubated with 5uM Rhodamine 123 for 20 mins. Then the cells were washed, harvested, pelleted, and resuspended in PBS and was used to perform flow cytometry (Cyflow® Space Partec, IL, USA) (excitation-488nm, emission-525 nm) (Scaduto & Grotyohann, 1999).

### 11. Evaluation of mitochondrial mass

Treated cells were incubated with 5µM NAO for 30 mins. Then the cells were washed, harvested, pelleted, and resuspended in PBS and was used to perform flow cytometry (Cyflow® Space Partec, IL, USA) (excitation-488nm, emission-525 nm) (Das et al., 2020).

### 12. Evaluation of mitochondrial oxidative stress

Oxidative stress in mitochondria was assessed using the dye MitoSOX red as mentioned by the manufacturer and the fluorescent intensity (excitation-510nm, emission-580 nm) was measured by flow cytometry (Cyflow® Space Partec, IL, USA).

### 13. Statistical analysis

Data are represented as mean ± SD for the experiments mentioned. Each of the experiments performed has been repeated at least thrice. The statistical significance between the experimental groups was calculated by one-way ANOVA followed by Dunnett’s multiple comparison test or by unpaired t test using GraphPad InStat software (GraphPad Software, La Jolla, CA,USA).

## RESULTS

### 1. Analysis of cell viability by MTT assay

MTT assays conducted to investigate the effect of Pb(NO_3_)_2_ on keratinocytes, mitochondrial dehydrogenases in viable cells, when treated with MTT convert the soluble MTT dye in the cells to an insoluble purple formazan product which is solubilized in DMSO and whose absorbance is directly proportional to the number of viable cells. Cells treated with a concentration of lead from 50-800 µM for 72 hrs showed a decrease in cell viability with an increase in Lead conc. compared to control and the IC_50_ concentration of Pb(NO_3_)_2_ in HaCaT cells were determined to be 385.7 µM (Fig. 1a).

**Fig. 1.**
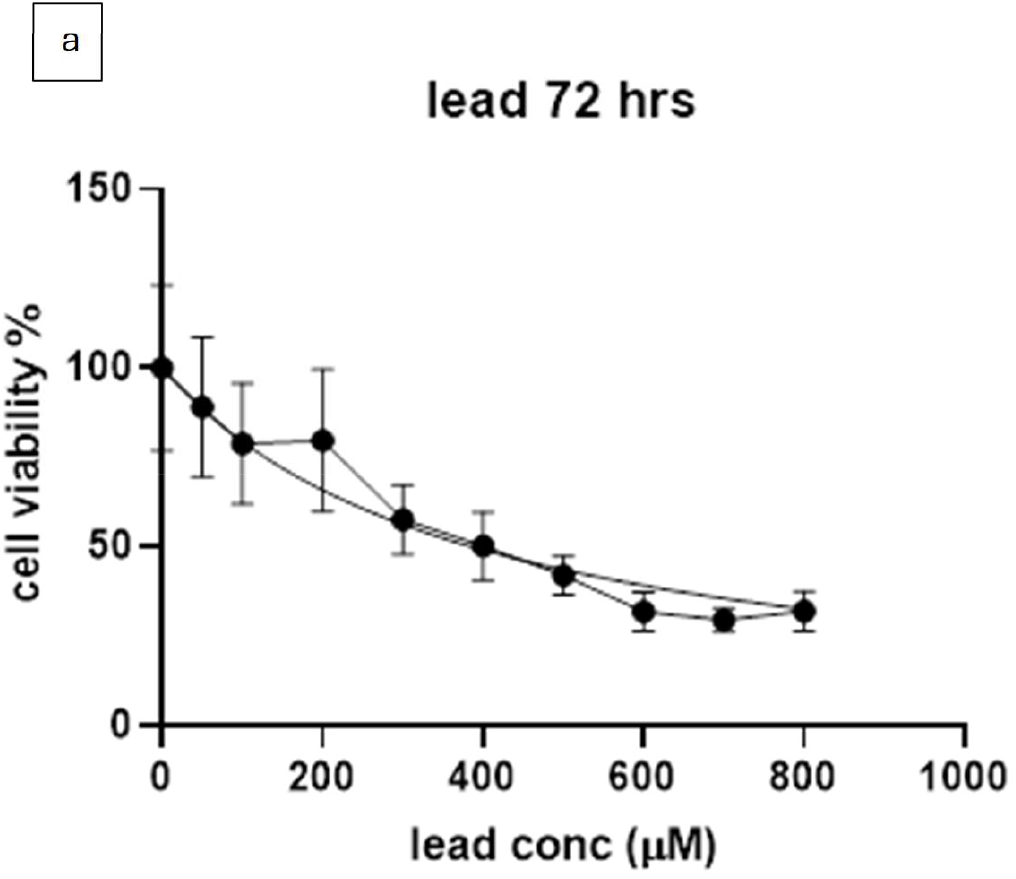
(a) Results of MTT assay with Pb(NO_3_)_2_ conc. taken from 50 µM -800 µM.

### 2. Clonogenic assay

The clonogenic assay revealed that the lead treated group had a much lower chance of creating a colony compared to control (p<0.05). While the Lead group only had a surviving fraction of 0.71 the control group had a surviving fraction of 1.00 (Fig. 2a).

**Fig. 2.**
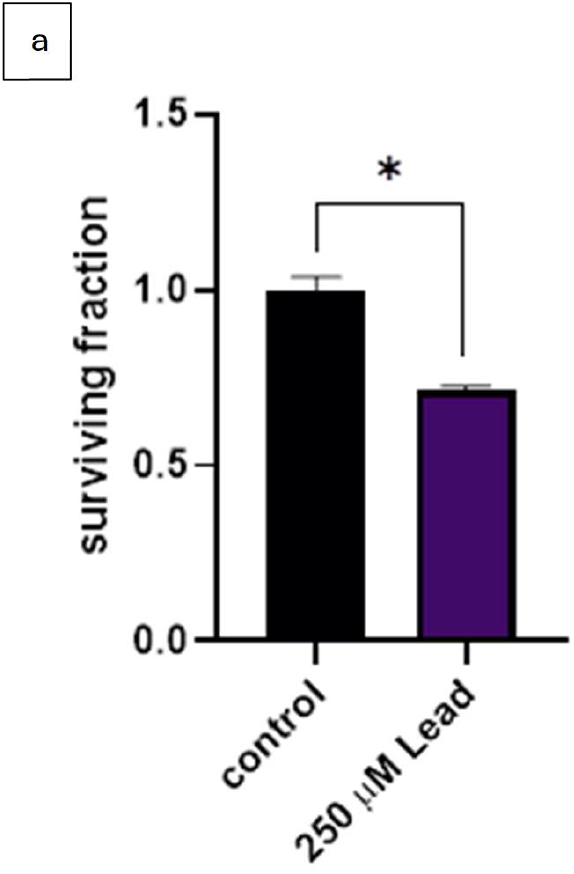
(a) Represents the surviving fraction of clonogenic assay. (*P < 0.05 when compared with control)

### 3. Micronucleus assay

The micronucleus assay was done by adding cytochalasin B to treated cells, which allowed the cells to undergo cell division but not cytokinesis due to disruption of actin filaments, which causes formation of binucleated cells, some of which contain a micronucleus. The number of BNC with MN was counted from 1000 BNC and the Lead treated group showed a significant increase in the amount of BNC with MN when compared to control (p<0.001). The Lead treated group had around 21 MN/1000 BNC, whereas the control group had only around 9.33 MN/1000 BNC (Fig. 3a).

**Fig. 3.**
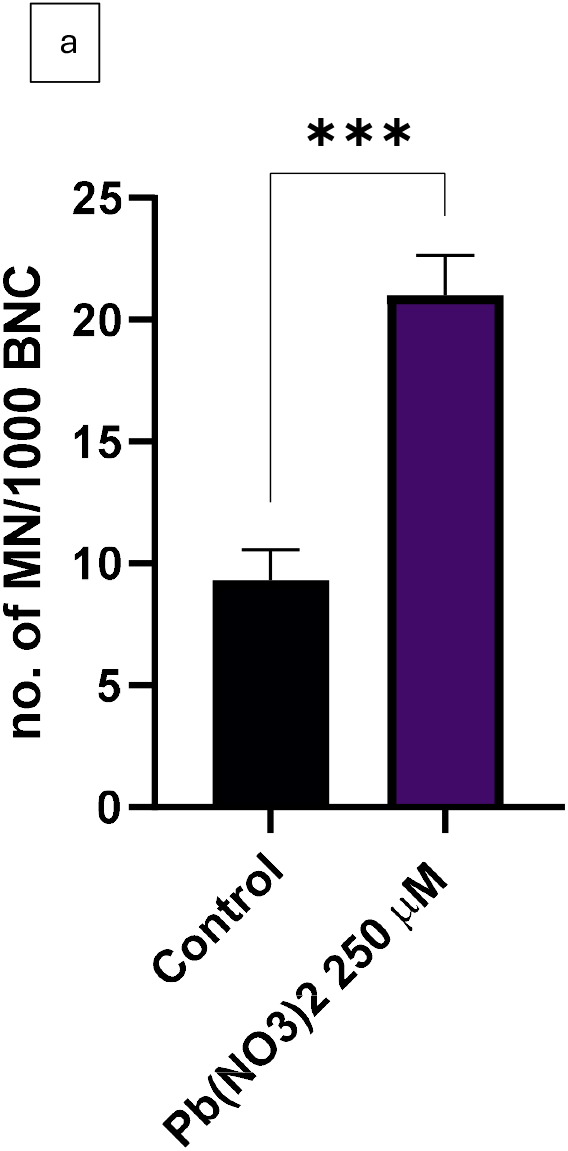
(a) Represents the no. of binucleate cells with micronucleus per 1000 binucleate cells (***P < 0.001 when compared with control).

### 4. Comet Assay

Alkaline single cell gel electrophoresis was done to assess the olive tail movement which is the product of DNA tail length and % of DNA in the tail. The olive tail movement of the lead treated group showed a significant increase compared to control (p<0.05). The Lead treated group had an average OTM of 10.62 µm, whereas the control group only had an average OTM of 4.34 µm (Fig. 4a).

**Fig. 4.**
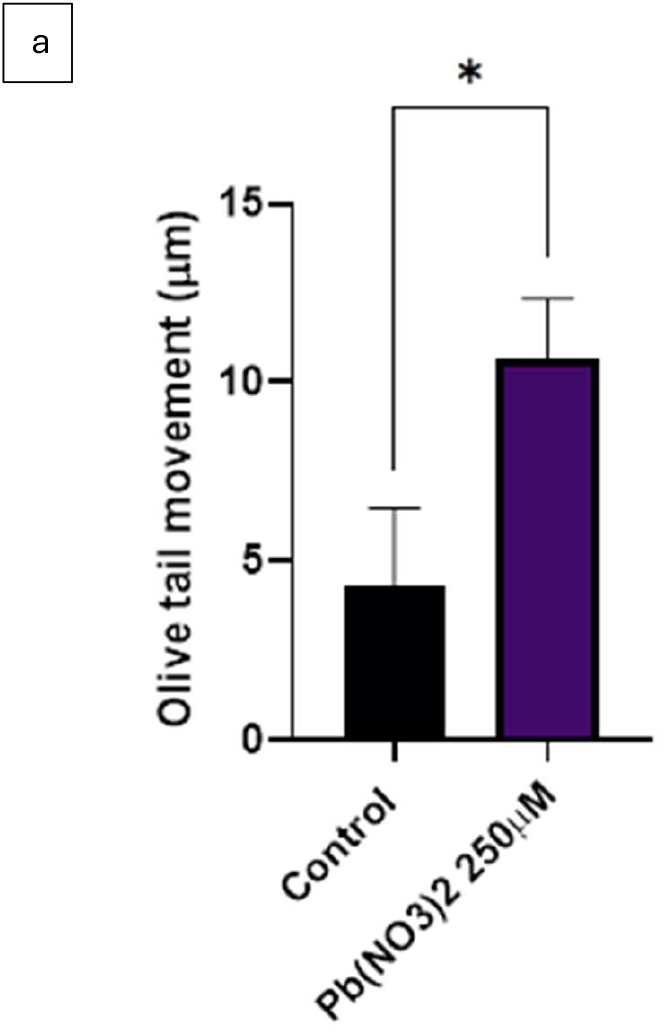
(a) Represents the change in olive tail movement of lead treated group compared to control, when alkaline single cell gel electrophoresis was conducted to assess DNA damage. (*P < 0.05 when compared with control).

### 5. Intracellular ROS production

Production of intracellular ROS was measured by DCF-DA which enters the cell and is converted to DCFH which is then oxidized to a fluorescent product called DCF by ROS (Bai et al., 1999), whose fluorescent intensity is measured. The lead treated group was found to have a higher amount of intracellular ROS as compared to control, although no significant difference was found. Mean fluorescent intensity (MFI) was calculated to be 13.59 for the control and 17.04 for the lead treated group (Fig. 5a).

**Fig. 5.**
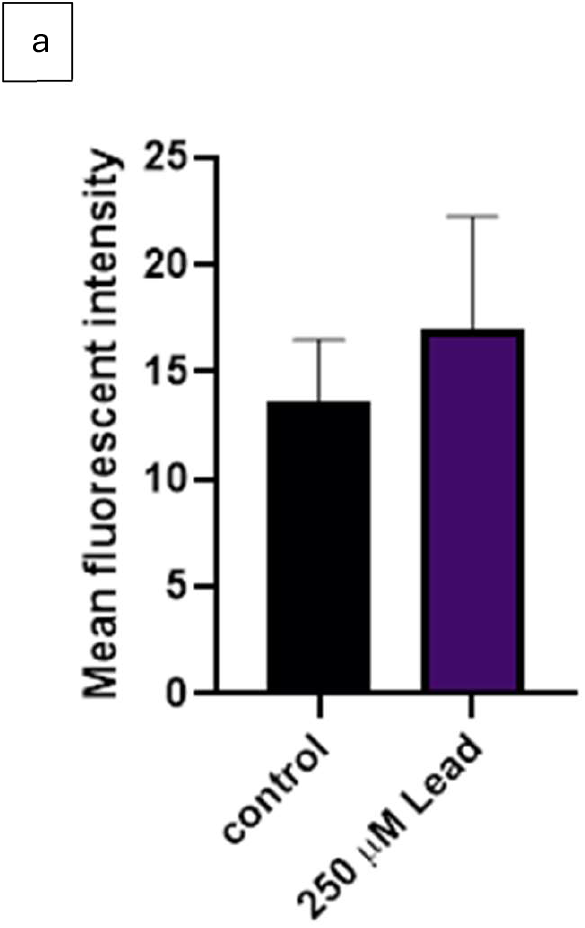
(a) Represents the change in mean fluorescent intensity of lead treated group compared to control, when treated with DCF-DA to assess intracellular ROS production.

### 6. Relative protein thiol content

Relative protein thiol content was estimated using DTNB, which reacts with thiol groups in proteins and forms a yellow-colored compound called TNB whose absorbance is measured. Cells treated with Pb(NO_3_)_2_ for 72 hrs showed a significantly lower amount of protein thiol content compared to control (p<0.0001), the lead treated group only had 46% of protein thiol compared to the control group (Fig. 6a).

**Fig. 6.**
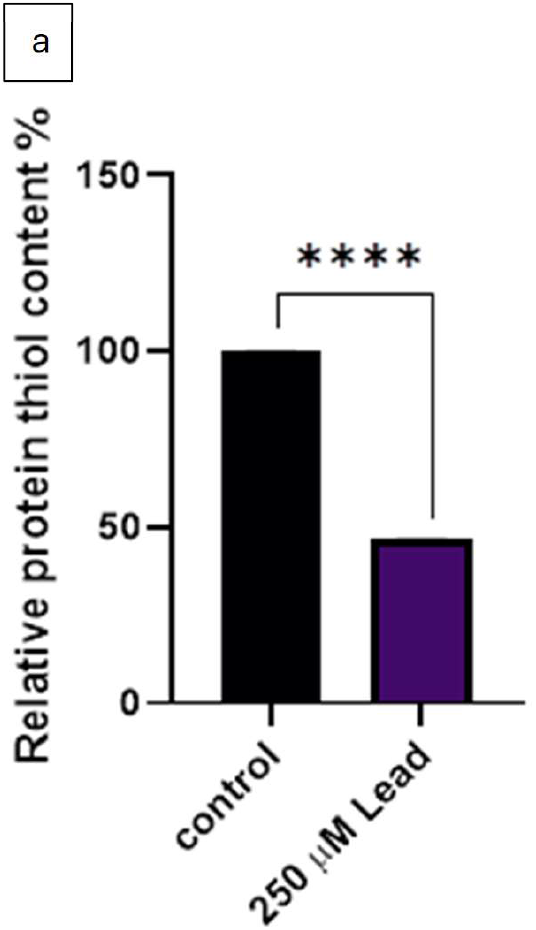
(a) Represents the relative protein thiol content of lead treated group compared to control. (***P < 0.001 when compared with control).

### 7. Mitochondrial membrane potential

Mitochondrial membrane potential was assessed using R123, which is concentrated inside the mitochondria as it is lipophilic and cationic and is retained within the mitochondria depending upon the membrane potential, and it was found that the lead treated group had an increased mitochondrial membrane potential when compared to control (p>0.05). The MFI was calculated to be 31.39 for the control and 36.59 for the lead treated group (Fig. 7a).

**Fig. 7.**
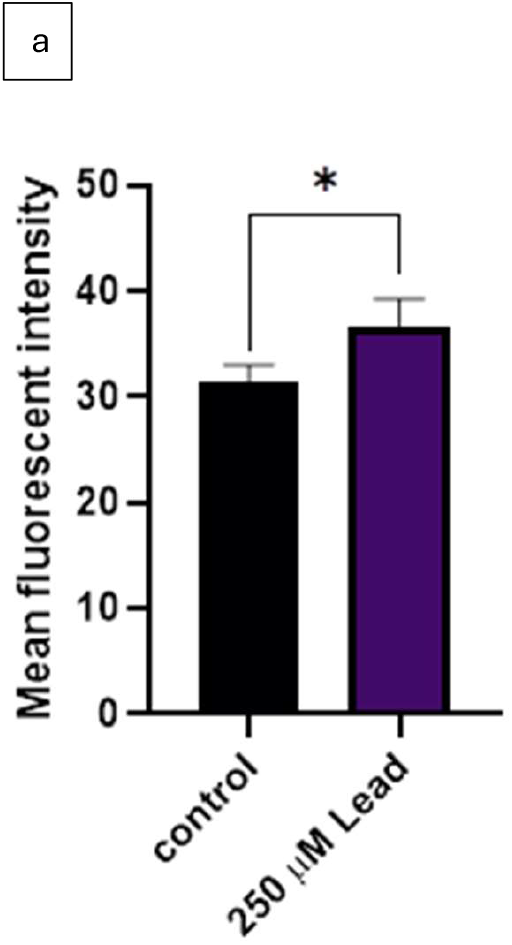
(a) Represents the change in mean fluorescent intensity of lead treated group compared to control, when treated with R123 to assess mitochondrial membrane potential. (*P < 0.05 when compared with control).

### 8. Mitochondrial Mass estimation

Mitochondrial mass was estimated using NAO which selectively binds to cardiolipin which is lipid found in the mitochondrial cell membrane, the fluorescent intensity is directly proportional to the amount of cardiolipin which corresponds to the mitochondrial mass. The flow cytometry analysis showed that the lead treated group had an increased mitochondrial mass compared to control (p>0.01). The MFI was calculated to be 46.61 for the control and 57.82 for the lead treated group (Fig. 8a).

**Fig. 8.**
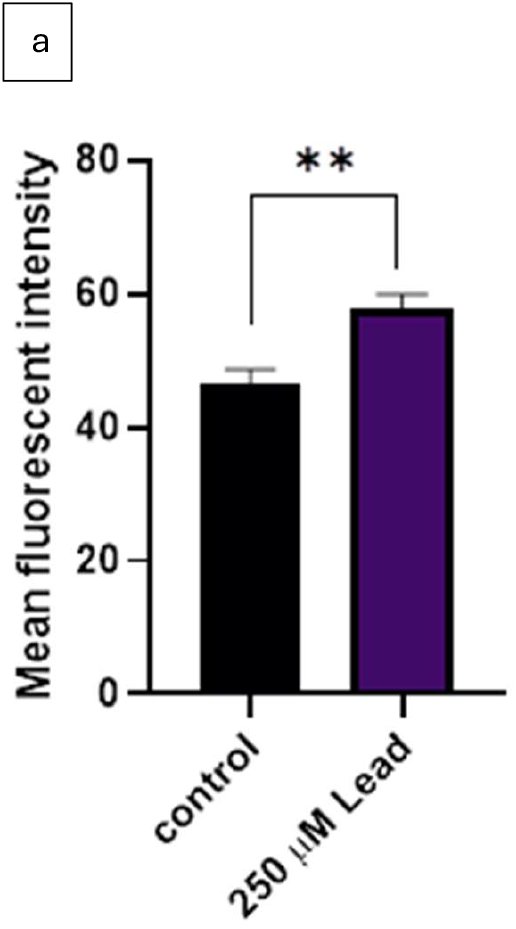
(a) Represents the change in mean fluorescent intensity of lead treated group compared to control, when treated with NAO to estimates mitochondrial mass. (**P < 0.01 when compared with control).

**Fig. 9.**
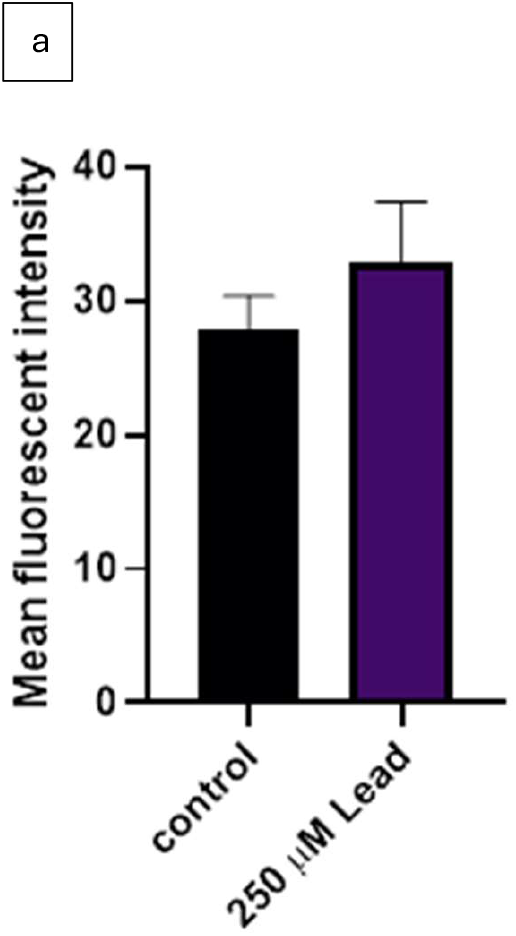
(a) Represents the change in mean fluorescent intensity of lead treated group compared to control, when treated with MitoSOX red dye to assess mitochondrial superoxide production.

### 9. Mitochondrial Superoxide production

The production of Mitochondrial Superoxide was measured using MitoSOX red dye, which selectively enters the mitochondria and is converted to its fluorescent product by mitochondrial superoxide. and the Lead treated group was found to have a higher amount of mitochondrial superoxide as compared to control, although no significant difference was found. The MFI was calculated to be 27.90 for the control and 32.78 for the lead treated group (Fig. 13a).

## DISCUSSION

The manifestations of the toxic effects of Lead, though often overlooked in research, are a very important marker for lead poisoning in clinical settings. The assays reported in this paper further prove the cytotoxic, genotoxic, effects of lead on keratinocytes as well as increased oxidative stress and altered mitochondrial function. This study found that Pb(NO_3_)_2_ caused a decrease in cell viability as concentration increased, the IC50 concentration was found to be 385.7 µM, a lower concentration of Pb(NO_3_)_2_, 250 µM which was approximately the IC30 concentration, was used to conduct rest of the assays. The clonogenic assay confirmed the results of MTT assay and showed a decrease in the percentage of surviving fraction (0.7133(L), 1.00(C)). The genotoxic assays also showed a similar trend where the lead group had a higher level of genotoxicity. The micronucleus assay showed a significant in the number of MNBNC (21) compared to control (9.33). And the comet assay confirmed those results showing a significant increase in the OTM (10.62 µm(L), 4.34 µm(C)) The assays conducted to show induced oxidative stress such as intracellular ROS and protein thiol content showed an increased intracellular ROS (MFI=17.04(L), 13.59(C)) although no significant difference was found, and a decreased protein thiol content (46%) for the lead treated group when compared to control. The mitochondrial superoxide levels were also higher in the lead treated group (MFI=32.78 (L), 27.90 (C)) but no significant difference was found. The mitochondrial mass and membrane potential were found to be higher in lead treatment compared to control (MFI= 57.82(L), 46.61(C)), (MFI=36.59(L), 31.39(C)) which does not concur with previously published data (Bandaru et al., 2023; Liu et al., 2014), the increase in mass may be due to an increase in mitochondrial fission which was suggested by Das and coworkers (Das et al., 2020), which maybe to maintain the energy requirement of the cell once exposed to lead, by increasing the total amount of mitochondria, cells can ensure that they still have enough healthy mitochondria to meet their energy needs, even while they are removing damaged ones, which could contribute to an increased cardiolipin content, although no assays were done to assess the credibility of this statement. The increase in fluorescence intensity, in case of R123 which may be due to hyperpolarization of the mitochondrial membrane which is involved in many cell death pathways and is known to be an early marker of apoptosis (Leal et al., 2015), ROS production has also been found to lead to hyperpolarization (Fedoseeva et al., 2017) in lead treated group or maybe due to quenching of the dye in the mitochondria of control group, or it could be due to the increased amount of mitochondria present in the treatment group as shown by the mitochondrial mass although the latter two statements are just speculations.

Conducting in vivo studies will be essential to validate the findings observed in vitro and gain insights into the systemic effects of lead exposure. Employing a diverse range of cell lines, including primary cells, will be crucial to corroborate results and enhance their applicability across various biological contexts. There is also a need to study the effects of long-term, low-dose lead exposure is imperative to simulate chronic environmental or occupational scenarios accurately. Deeper mechanistic studies are also needed to elucidate the molecular pathways through which lead induces oxidative stress and mitochondrial dysfunction, which could lead to identification of novel targets for therapeutic intervention.

This study sets the stage for further analysis of the toxic effects of lead on skin, which would greatly increase the speed and efficiency of lead poisoning diagnoses in clinical settings, enhance safety standards and protective measures in various industries by elucidating the mechanisms of dermal absorption and informing the development of more effective protective gear and skincare products. Such insights could also shape public health policies, particularly in safeguarding vulnerable populations, and guide the formulation of medical treatments to mitigate the health impacts of lead exposure. A deeper understanding of lead’s dermal toxicity could drive more stringent testing and regulation of consumer products, thereby ensuring greater safety in everyday items such as cosmetics, jewelry, and toys.

## CONCLUSION

Lead nitrate was found to be toxic to HaCaT cells, and the IC_50_ concentration was determined to be 385.7 µM at 72 hrs. Lead nitrate at 250 µM was also found to decrease the clonogenic potential, produce an increased number of binucleated cells with a micronucleus, reduce the thiol protein content, increase the intracellular and mitochondrial ROS levels although this study did not find a significant increase, increase the mitochondrial mass and mitochondrial membrane potential, which could be due to increased mitochondrial fragmentation due to mitochondrial fission. Which shows that Lead nitrate does indeed show cytotoxic, genotoxic effects as well as increase oxidative stress and alter mitochondrial functions.

## Declaration of competing interest

The authors declare that they have no known competing financial interests or personal relationships thsat could have appeared to influence the work reported in this paper.

### Declaration of Generative AI and AI-assisted technologies in the writing process

During the preparation of this work the authors used ChatGPT to assist with refining the text. After using this tool, the authors reviewed and edited the content as needed and takes full responsibility for the content of the publication.

## Acknowledgments

The authors would like to thank the Manipal School of Life Sciences and Manipal Academy of Higher Education (MAHE) for the infrastructure and facility.

